# Very few sites can reshape a phylogenetic tree

**DOI:** 10.1101/413518

**Authors:** Warren R. Francis, Donald E. Canfield

## Abstract

The history of animal evolution, and the relative placement of extant animal phyla in this history is, in principle, testable from phylogenies derived from molecular sequence data. Though datasets have increased in size and quality in the past years, the contribution of individual genes (and ultimately amino acid sites) to the final phylogeny is unequal across genes. Here we demonstrate that by removing a small fraction of sites that strongly favor one topology, it is possible produce a highly-supported tree of an alternate topology. We explore this approach using a dataset for animal phylogeny, and create a highly-supported tree with a monophyletic group of sponges and ctenophores, a topology not usually recovered. As nearly all gene sets are neither standardized nor representative of the entire genome, we conclude that there are only two possible ways to remedy this problem. One solution would need to use a fixed set of genes, which though not representative, is at least standardized. The other would be to construct phylogenies using all genes, thus limiting analysis to species with sequenced genomes.

## Introduction

It has been over a decade since Rokas et al. (2005) noted substantial challenges in reconciling molecular phylogeny of metazoans, particularly with respect to deep nodes. In an early attempt to apply molecular sequence data to bilaterian evolutionary relationships, Dunn et al. (2008) had the surprising finding that ctenophores (comb jellies) emerged as the sister-group to the rest of metazoans (hereafter called Ctenophora-sister), contrary to the classically-held view that sponges were sister-group to all other animals (the hypothesis called Porifera-sister). A number of papers followed arguing both for and against each of these topologies (Philippe et al. 2009, Ryan et al. 2013, Whelan et al. 2015, Pisani et al. 2015, Simion et al. 2017, Whelan et al. 2017). Thus, despite over a decade of work, the deep branches of the animal tree remain unresolved.

The choice of genes used in phylogenetic reconstruction may have a substantial effect on the final tree. Shen et al. (2017) have shown that for most controversial nodes, some genes have very strong phylogenetic signals while other genes contain essentially none. While Shen et al. (2017) made some suggestions about how to resolve recalcitrant nodes, their method highlighted a potential risk of “stacking the deck” and generating a biased tree topology by selecting a set of genes that skew towards one topology. Here we demonstrate that with the removal of only 1.7% of sites, we can generate a tree with an alternate topology of metazoan phylogeny. We then discuss the overall lack of scrutiny on sitewise filtering strategies and suggest that substantial biases can be introduced.

## Methods

### Datasets and processing

We re-analylzed dataset 16 from Whelan et al. (2015), the same dataset used in the re-analysis by Pisani et al. (2015) and by Shen et al. (2017). This dataset was a filtered version of the main dataset used by Whelan et al. (2015), wherein genes and taxa with high long-branch scores were removed, and from that, the slowest-evolving half of the genes were analyzed.

Sitewise likelihood calculations were generated using the method of Shen et al. (2017), with one difference. Briefly, this is a four-branch resolution problem, whereby the method takes three fixed trees and analyzes the likelihood at each site given the three possibilities (Figure 1). Using the program RAxML, this is done with the option -f G. The likelihood values for each site for each tree are then directly compared, where the least negative means the most likely. However, in Shen et al. (2017), the strength of the site for each topology (dlnL) was calculated as the average of the absolute value of the three differences. Such approach would overestimate the strength of sites where one topology was substantially weaker (i.e. less likely) than the other two. Thus, we defined the strength of a site as the values of the maximum likelihood topology minus the score of the second best topology. Here “strong sites” are defined as sites where the absolute difference in log likelihood is greater or equal to 0.5, the same threshold used by Shen et al. The vast majority of sites have differences in likelihood values that are close to zero (appx. 98% of sites), thus a dlnL score of 0.5 represents roughly 3 standard deviations above the mean.

**Figure 1:**
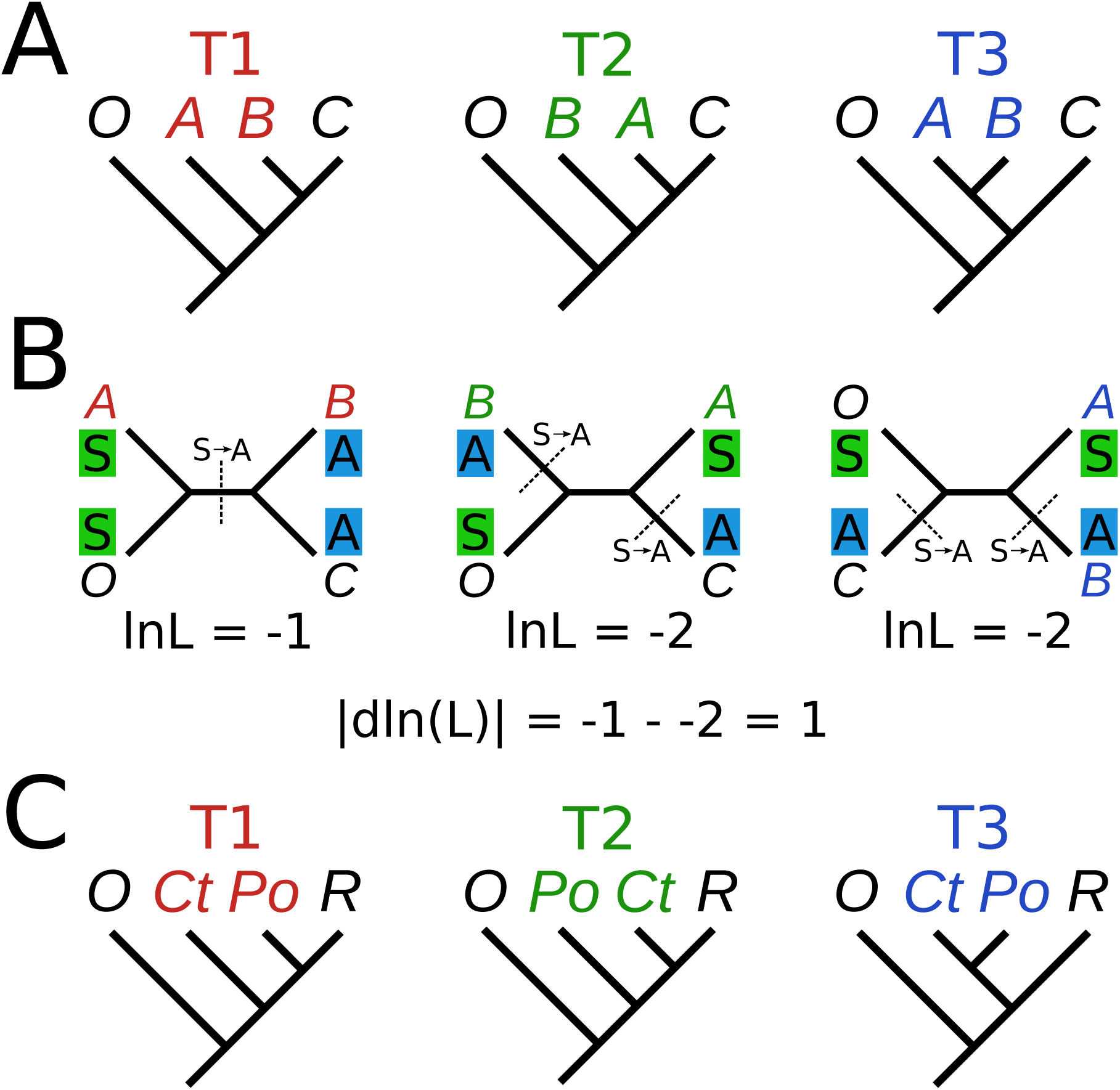
Schematic of analysis: (A) Three fixed trees differ by the position of groups A and B, relative to group C and outgroup O. (B) Sites in the alignment either show 1 or 2 substitutions, depending on which tree is used. The substitutions do not have direction in time-reversible models, so the transition applies in either direction across the dotted lines. In this hypothetical example, the dln(L) between the maximum (−1) and median (−2) would be 1, indicating a strong site favoring T1. In this case, while T1 has the maximum likelihood, it is also the most parsimonious. (C) Concretely, in our study, T1 was the Ctenophora-sister hypothesis, T2 was the Porifera-sister hypothesis, and T3 was the paranimalia hypothesis. ‘Ct’ and ‘Po’ indicate ctenophores and sponges, respectively, while ‘O’ indicates non-metazoan outgroups (other opisthokonts) and ‘R’ indicates the rest of animals.

To generate our experimental dataset (called the “weak” dataset), we started with the site-wise likelihood scores from Shen et al. (2017) for dataset D16 of Whelan et al. (2015), which were reformatted to a tabular file using a Python script sitewise_ ll _ to _ columns.py. This was then used as the input for another script sitewise _ get _ strong _ sites_ 2tree.py that calculated strong sites based on the first two trees, Ctenophora-sister and Porifera-sister, and removed sites with dlnL greater or equal to 0.5 that favored either of the two topologies, but not those supporting the third topology, the monophyly of sponges and ctenophores. This procedure removed 414 sites out of the total 23676 sites, only 1.7% (for comparison, human and zebrafish are 14% different in this dataset.) These scripts can be found at the Github repository https://github.com/wrf/ pdbcolor/tree/master/sitewise_scripts.

### Phylogenetics

We generated phylogenetic trees using RAxML v8.2.11 (Stamatakis 2014) using the PROTGAMMALGF model and 100 bootstrap replicates with the “rapid boostrap” option (-f a). The same dataset was run in a Bayesian framework with Phylobayes-MPI v1.8 using the CAT-GTR model (Lartillot et al. 2007). Two chains were run in parallel for approximately 1000 cycles and otherwise default parameters. Trees and run data can be found at the online repository https://bitbucket.org/ wrf/paranimalia-sites.

### Comparison across datasets

We compared the extent of alignment trimming and sitewise coverage across several phylogenetic datasets from previously published studies (see Table 1). For calculation of the trimmed fraction for each protein, we used the length of the alignment relative to the human reference protein. Because human proteins were not included in the Philippe 2009 or Ryan 2013 EST datasets, the human orthologs needed to be identified for each gene.

**Table 1:** Phylogenomic data sources

| Dataset name | Genes | Taxa | Sites | Average coverage (by gene) | Average coverage (by site) | Human proteins | Total kept fraction after trimming | Reference |
| --- | --- | --- | --- | --- | --- | --- | --- | --- |
| Philippe 2009 | 128 | 55 | 30257 | 82% | 73.1% | 0 (128) | 86.0% | (Philippe et al. 2009) |
| Ryan 2013 EST | 406 | 70 | 88384 | 50 | 41.6 | 0 (396) | 62.5 | (Ryan et al. 2013) |
| Whelan 2015 D1 | 251 | 76 | 81006 | 75 | 59.6 | 248 | 56.6 | (Whelan et al. 2015) |
| Borowiec 2015 | 1080 | 36 | 384981 | 87 | 75.8 | 1056 | 64.7 | (Borowiec et al. 2015) |
| Cannon 2016 | 212 | 78 | 44896 | 80 | 69.0 | 212 | 46.1 | (Cannon et al. 2016) |
| Simion 2017 | 1719 | 97 | 401632 | 74 | 60.7 | 1499 | 40.0 | (Simion et al. 2017) |

We developed a pipeline to identify genes from an existing alignment in additional species, called _ add _ taxa _ to _ align.py. This pipeline makes use of hmmbuild and hmmsearch from the HMMER package v3.1b2 (Eddy 2011) and the alignment program MAFFT v7.313 (Katoh et al. 2017). Briefly, for each gene in a supermatrix, a hidden Markov model is generated using hmmbuild, and this is used as the query for hmmsearch to search within a file of proteins from the new species. The results are filtered by multiple heuristics, and the best sequence is added to the existing alignment using MAFFT, with the --addlong option.

This script and related instructions are available at the github repository: https://github.com/wrf/supermatrix.

## Results

### Paranimalia is recovered regardless of model

By removing the “strong” sites from the supermatrix alignment, we then generated two phylogenetic trees using two programs, RAxML (using the model PROTGAMMALGF) and phylobayes (under the model CAT-GTR) to assess the impact on the final tree.

As expected, both trees strongly supported monophyly of ctenophores and sponges, (bootstrap:94; PP:1.0; Figure 2), hereafter called “paranimalia”. This confirms that the sites removed contained the majority of phylogenetic information in support of Ctenophora-sister or Porifera-sister. Although the matrix contained distant outgroups (fungi, as well as choanoflagellates) (Philippe et al. 2009, Pisani et al. 2015), Ctenophora-sister was not recovered in either tree, indicating that either any long-branch attraction artifacts are weaker than the intrinsic signal in the sites, or the sites that support Ctenophora-sister are those subject to the proposed “long branch attraction”.

**Figure 2:**
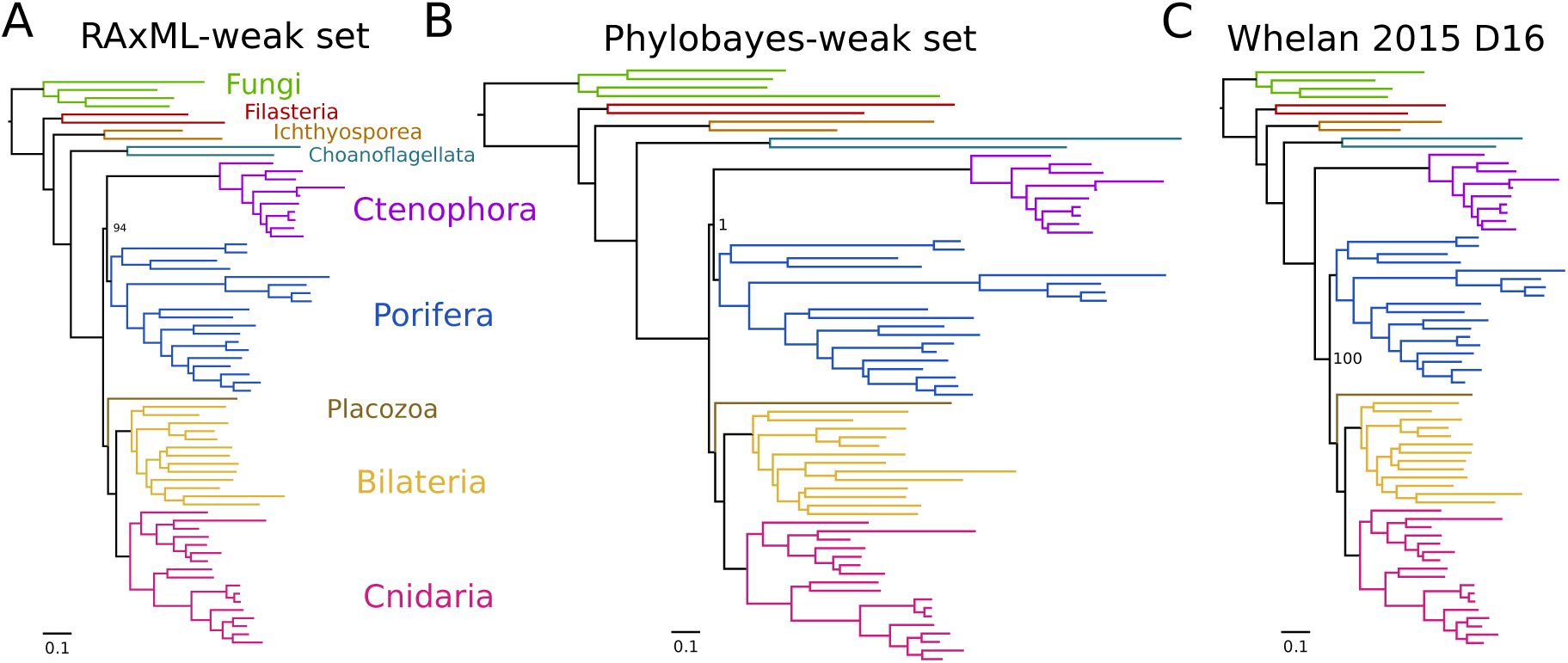
Overview of phylogenetic trees: (A) Tree from RAxML (B) Tree from Phylobayes with CAT-GTR. Note the scale bars are the same but the phylobayes tree is substantially longer, likely due to increased substitutions predicted from the CAT model. (C) Original tree from Whelan et al. (2015) dataset 16, showing that, other than ctenophores and sponges, nearly all bipartitions are exactly the same before and after our processing. Most support values are removed for clarity.

### Few topological differences are found

The internal topology of nearly all phyla remains the same (Figures 3 and 4), despite changing the position of the nodes for ctenophora and porifera, suggesting that sites providing information for each bipartition are mostly independent. One obvious inconsistency is the placement of Ichthyosporea and Filasterea relative to dataset 10 by Whelan et al. (2015), as these two groups are swapped (see both Figures 3 and 4).

**Figure 3:**
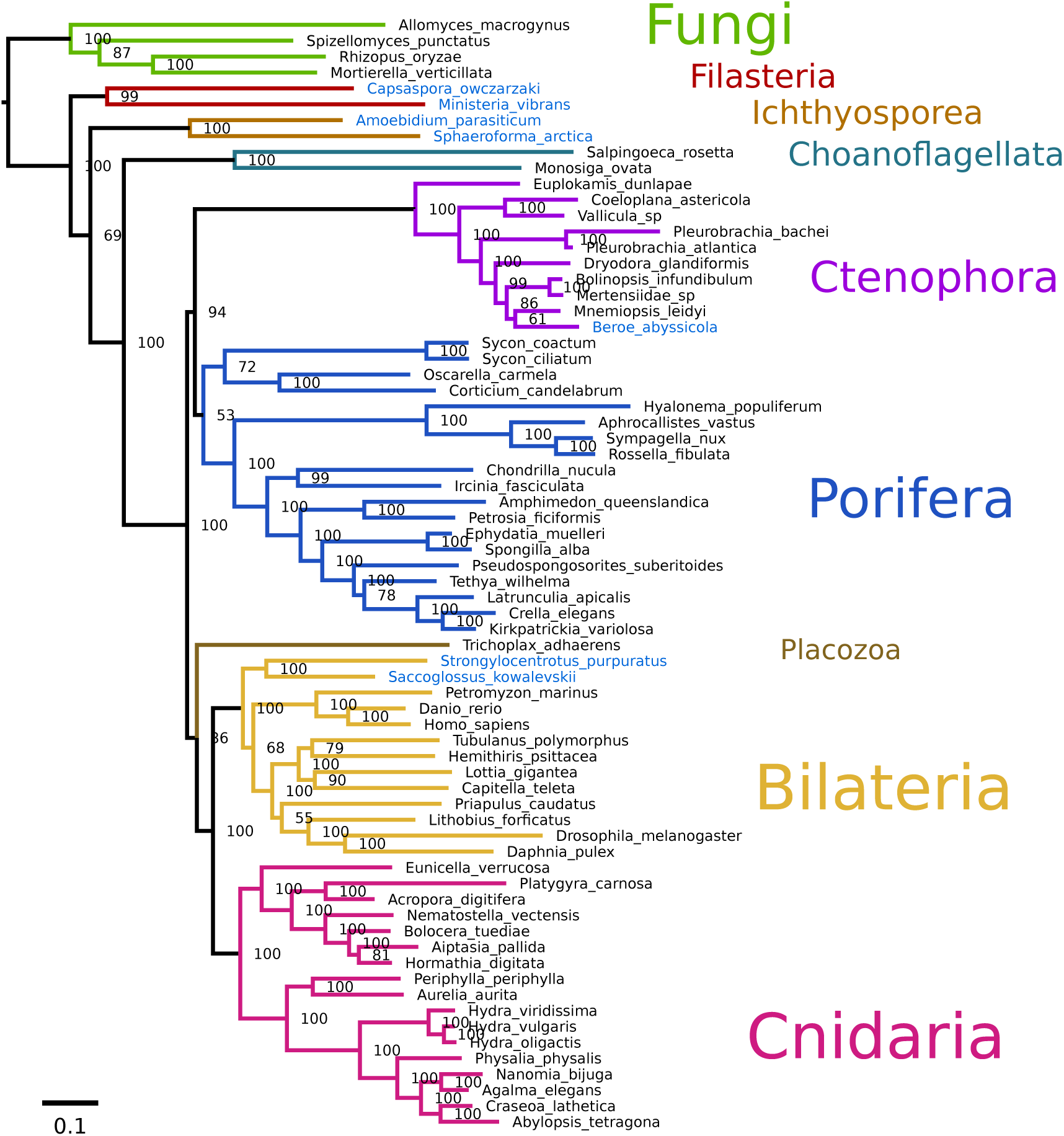
RAxML tree: Tree of the weak dataset from RAxML using the PROTGAMMALGF model. Taxa highlighted in red are moved relative to the original dataset 16 by Whelan et al. (2015). Taxa highlighted in blue are moved relative to dataset 10 from the same study.

**Figure 4:**
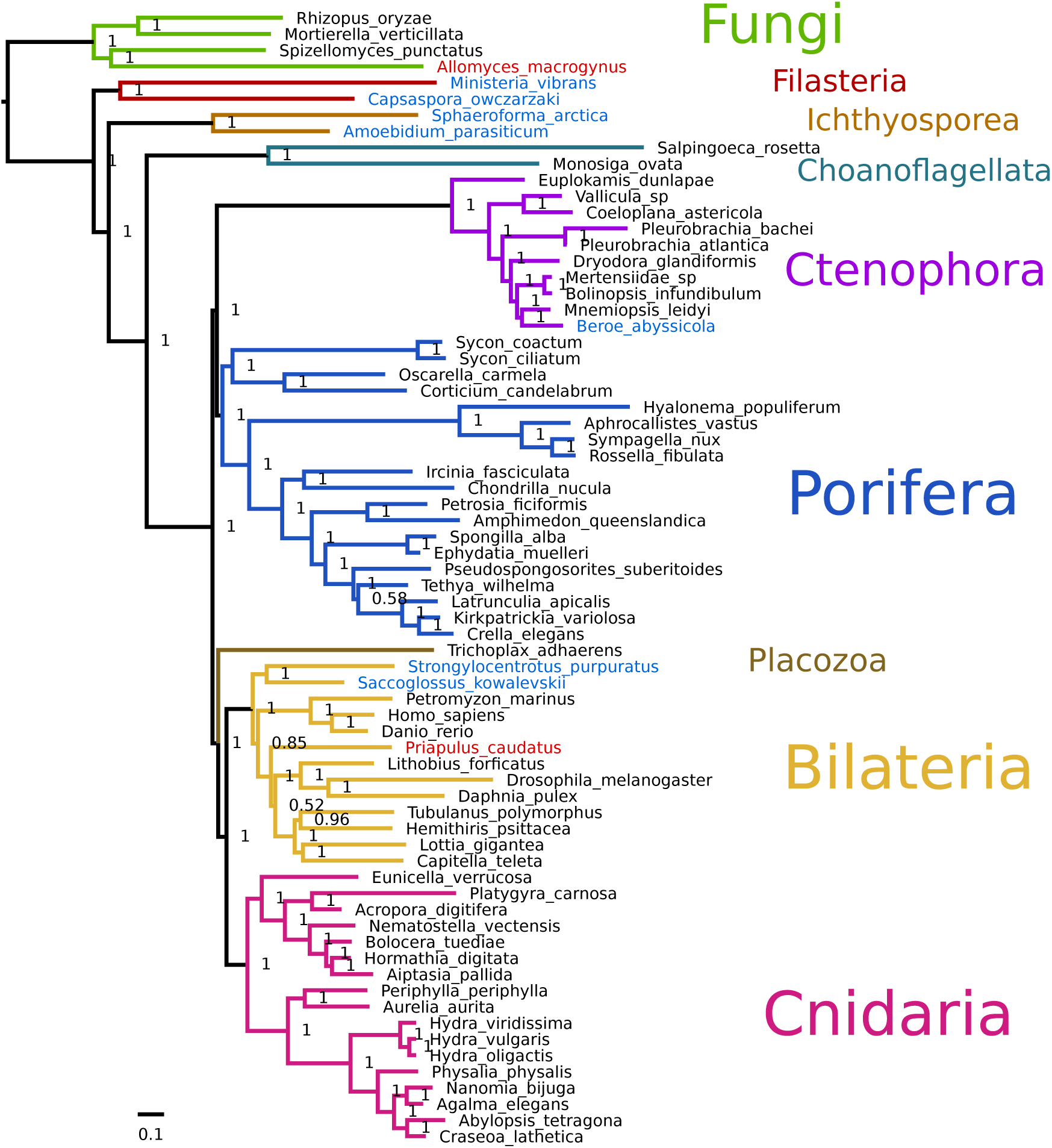
Phylobayes tree: Tree of the weak dataset from phylobayes using the CAT-GTR model. Taxa highlighted in red are moved relative to the original dataset 16 by Whelan et al. (2015). Taxa highlighted in blue are moved relative to dataset 10 from the same study.

For both analyses, Ambulacraria was recovered as sister to Bilaterians (71 bootstrap, PP 0.85), indicating paraphyletic deuterostomes. This topology was also recovered by Whelan et al. (2015) with their dataset 16, but not with dataset 10, which was used for the main figure (Figure 3 in Whelan et al. (2015)). This position of Ambulacraria was also found by Simion et al. (2017) after substantial trimming of the dataset, whereupon 70% of “heteropecillious” sites were removed (Roure and Philippe 2011). Cannon et al. (2016) found this tree position occupied by the Xenacoelamorpha; this group includes the genus *Xenoturbella*, which was often recovered as sister to Ambulacraria within Deuterostomes (Philippe et al. 2009; 2011). The recovery of Ambulacraria as sister to Bilaterians may indicate a relationship between heteropecilly (lineage-specific transitions or substitution matrices) and strong sites in a maximum likelihood framework. In other words, lineage-specific changes in proteins may be a major source of phylogenetic information.

Other small differences are evident (Figures 3 and 4), such as the placement of the ctenophore *Beroe abyssicola* relative to *Mnemiopsis leidyi* (PP:1). Another is the placement of *Priapulus caudatus* as sister to protostomes, instead of just arthropods (PP:0.52).

### Most datasets are heavily trimmed

In our weak dataset, only 1.7% of sites had been removed, albeit the original dataset had already been trimmed in the D16 by an average of 30% per protein, compared to the reference proteins. While such trimming strategies make sense for highly repetitive regions that cannot be aligned, one study found that nearly all programs will overtrim, resulting in an overall less-supported phylogeny than if nothing were done at all (Tan et al. 2015). Even across these six studies in Figure 5 that we investigated, none of them include an unfiltered version for analysis, so the effect of these removed sites or domains is unknown.

**Figure 5:**
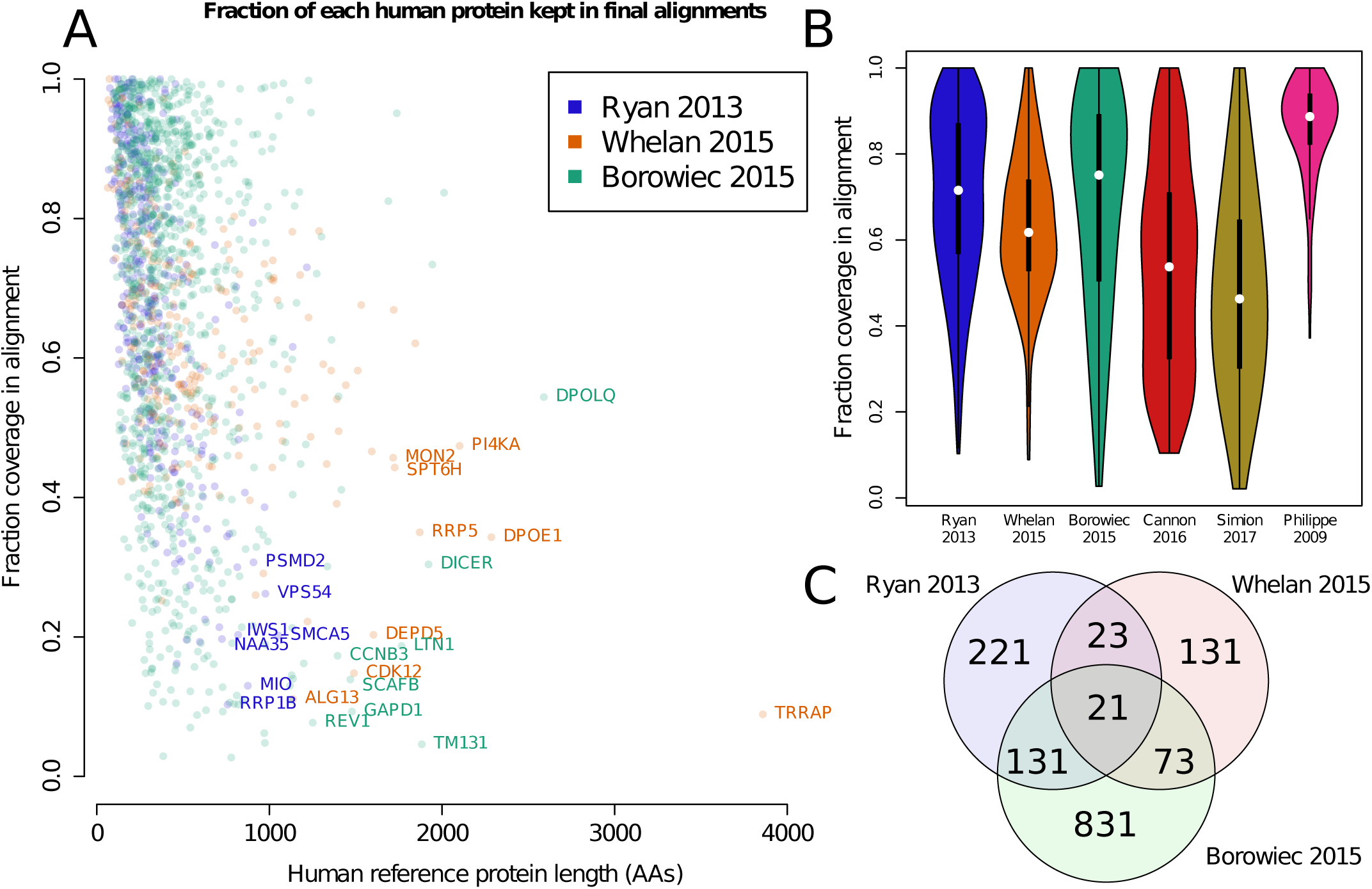
Fraction of the original protein in the final alignment: (A) Fraction of the original human protein that was used in the final alignment, meaning after any trimming steps, for the three metazoan datasets used by Shen et al. (2017), that is, the datasets from Ryan et al. (2013), Whelan et al. (2015), and Borowiec et al. (2015). Certain large and highly-trimmed protein names (Uniprot IDs) are indicated. (B) Violin plots showing the distribution of the kept fractions of the original proteins across the six datasets. Width is proportional to the normalized number of proteins with the coverage on the Y-axis. Simion 2017 was the least retained after trimming (average of 48%) while Philippe 2009 was the most (average of 86%). (C) Venn diagram of the proteins used in each study, out of 1431 total proteins across all three studies.

As there is typically no examination of which sites are filtered, it is easy to imagine the incidental removal of sites favoring a particular hypothesis, as we had specifically done in this study. The most trimmed study, (Simion et al. 2017) had removed over half of the amino acids of each protein on average compared to the reference proteins, from almost 1 million amino acids of total native proteins to an alignment of just over 400k amino acids (Table 1). As sites affecting deep nodes may account for only a fraction of 1% of all sites, exclusion of 60% of the original sites may affect deep nodes but not shallow nodes.

## Discussion

### Utility of sitewise filtering

Shen et al. (2017) had analysed the contributions of individual sites against trees with a fixed topology to discern which sites favor each tree. While this work did not resolve many of the controversial phylogenies, including that of animals, it did emphasize the importance of gene selection. Such a method is highly sensitive to taxon sampling, and likelihood scores can be calculated even in cases where it appears to be inappropriate, or makes little biological sense. For instance, in the Borowiec et al. (2015) dataset, there was only one ctenophore and one sponge, yet strong sites favoring Ctenophora-sister or Porifera-sister were still calculated even when the gene was absent for one or both of the two species. Potentially, genes where the ctenophore or sponge were absent should have been excluded. Therefore, in order for the results to be meaningful, essentially all sites should be occupied for all relevant taxa, in this case meaning all ctenophores, sponges, and outgroups.

### Other examination of bias in datasets

The work by Feuda et al. (2017) had attempted to examine the effects of strong sites as a function of substitution model. However, the “outlier-excluded” dataset used in their re-analysis was produced by removing outliers without considering the topology they favored and this site-selection methodology actually resulted in a dataset depleted in Ctenophora-sister favoring sites (all of the seven outliers favored Ctenophora-sister). A tree supporting Porifera-sister should therefore be expected from the analysis of this dataset as genes strongly supporting Ctenophora-sister were removed, but not those favoring Porifera-sister or any other systematic bias. Our results indicated that removal of sites favoring a specific topology (in our case, both Ctenophora-sister and Porifera-sister) can produce a highly supported tree favoring another topology for which sites were not removed (i.e. paranimalia).

### Resolution of metazoan phylogeny

Relationships among non-bilaterian animals historically placed sponges as sister-group to remaining animals, which agreed with a scheme of “complexity” coming from the Aristotelian chain-of-being; sponges are simple animals, and therefore should be placed at the root of the animal tree. Although by this logic, the morphologically simplest animals, placozoans, should therefore be the sister-group to all other animals. Ctenophores, on the other hand, have historically been placed in a group with cnidarians, called “coelenterata” or “radiata”, though detailed morphological analyses argued that every proposed synapomorphy of “coelenterata” was either uninformative or incorrect (Harbison 1985), indicating they were falsely united. The “paranimalia” hypothesis was only generated here as an example, but these two phyla (Porifera and Ctenophora) are united by some qualities, such as the absence of the HIF oxygen sensing pathway (Mills et al. 2018).

Animal molecular phylogeny remains controversial and technically challenging because differences in gene set (Nosenko et al. 2013), substitution model (Feuda et al. 2017), and missing data (Roure et al. 2013) have profound effects on the final tree. Other technical factors like introduction of newer versions of software make it practically infeasible to compare between datasets and results. In practice, this might require downloading or re-assembling the source data, finding orthologous genes across all species, filtering paralogs or incomplete transcripts, aligning, trimming, and finally generating the tree.

There is the additional semantic problem of how the results are described. There are almost an infinite number of possible hypotheses of metazoan phylogeny; most of these are unlikely, thus we may concern ourselves with a limited set of competing hypotheses of animal phylogeny, Ctenophora-sister and Porifera-sister. It is common to say there is “robust” support for a hypothesis in phylogenetics based purely on the bootstrap or posterior probability, but these two values do not reflect the fraction of sites favoring the two hypotheses of interest. Even considering the results of Shen et al. (2017) at face value, the only datasets that have reasonable coverage of ctenophores and sponges (meaning the sitewise likelihood is based on more than 1 taxon per phylum) are the Whelan et al. (2015) datasets. Of these, barely above 50% of all sites (both strong and weak sites) favor Ctenophora-sister. Shen et al. (2017) argued this was still sufficient support for the Ctenophora-sister hypothesis. However, this was without consideration of inherent noise in the data. For instance, sitewise likelihood values are calculated for all sites, including constant sites, and including sites with no ctenophores or no sponges. As weak sites are essentially phylogenetic noise, it would be more accurate to say this hypothesis is *slightly* or *marginally* favored, while 98% of sites do not affect this part of the tree.

Our results indicate that removal of a small fraction of sites (under 2%) can dramatically change the tree, and ultimately the hypotheses of animal evolution, yet many studies trim at least 40% of sites from the reference proteins (Figure 5). Thus, the resolution of the deep nodes of the tree, regardless of method or model, is extremely poor, and the statistical strategies to validate the approach (bootstrapping or posterior probability) do not reflect the true uncertainty of the data. Given the tenuous support for any of the topologies of animal phylogeny, it seems reasonable to say that we simply lack the information to resolve this, and should, at this time, defer on the null hypothesis that this node is still unresolved.

### What would make an unbiased set?

There are only two possibilities to have an unbiased set, whether deliberately or algorithmically. One would be a finite set of select genes that most or all species have and everyone agrees to use, such as mitochondrial proteins or ribosomal proteins. These may not be representative of the entire genome (potentially a bias in itself) but could at least be standardized. The other option would be to include all proteins, including those with multiple copies. Because of the difficulty in resolving species trees from multi-copy gene trees, algorithmic improvements may also be necessary. This may require that all species used in phylogenetic reconstructions have sequenced genomes to ensure that all genes are sampled, as bona fide gene losses cannot be identified with transcriptomes.

## Acknowledgements

WRF would like to thank SHD. Haddock, S. Vargas and M. Eitel for helpful comments. This work was supported by the Villum Foundation grant 16518 to DEC.

